# Tiara: Deep learning-based classification system for eukaryotic sequences

**DOI:** 10.1101/2021.02.08.430199

**Authors:** Michał Karlicki, Stanisław Antonowicz, Anna Karnkowska

## Abstract

**Motivation:** With a large number of metagenomic datasets becoming available, the eukaryotic metagenomics emerged as a new challenge. The proper classification of eukaryotic nuclear and organellar genomes is an essential step towards the better understanding of eukaryotic diversity.

**Results:** We developed Tiara, a deep-learning-based approach for identification of eukaryotic sequences in the metagenomic data sets. Its two-step classification process enables the classification of nuclear and organellar eukaryotic fractions and subsequently divides organellar sequences to plastidial and mitochondrial. Using test dataset, we have shown that Tiara performs similarly to EukRep for prokaryotes classification and outperformed it for eukaryotes classification with lower calculation time. Tiara is also the only available tool correctly classifying organellar sequences.

**Availability and implementation:** Tiara is implemented in python 3.8, available at https://github.com/ibe-uw/tiara and tested on Unix-based systems. It is released under an open-source MIT license and documentation is available at https://ibe-uw.github.io/tiara. Version 1.0.1 of Tiara has been used for all benchmarks.

## 1. Introduction

Microbial communities of unicellular eukaryotes (protists) and prokaryotes are an essential part of all ecosystems. Next to prokaryotes, protists are significant drivers in diverse nutrient cycling pathways. Autotrophic and mixotrophic protists fix carbon in aquatic environments, whereas heterotrophic protists catalyze nutrient cycling in aquatic and terrestrial ecosystems as selective consumers of bacteria and fungi (Caron *et al.*, 2009).

Metagenomic studies changed our understanding of the prokaryotic communities and allowed us to uncover their taxonomic and functional diversity in various environments (Sunagawa *et al.*, 2015; Almeida *et al.*, 2019). However, even though microeukaryotes are key components of microbial communities, their study lags behind the study of prokaryotes (Keeling and Campo 2017), and that is particularly true for the metagenomic studies. Until now, mainly metabarcoding (e.g. De Vargas *et al.*, 2015), metatranscriptomics (e.g. Salazar *et al.*, 2019), and single-cell genome sequencing (e.g. Strassert *et al.*, 2018) were used to explore the diversity of microbial eukaryotes. In contrast to these methods, the utilization of the metagenomic approaches was hampered by the complexity and size of eukaryotic genomes, as well as a limited number of reference databases allowing further taxonomical or functional annotation. With a few exceptions, such as phytoplankton (Delmont *et al.*, 2015; Duncan *et al.*, 2020) or human microbiome (Olm *et al.*, 2019) studies, eukaryotes were neglected in metagenomic studies. Only recently the metagenomic datasets from large sampling projects, such as the Tara Oceans expedition (Pesant *et al.*, 2015) or Ocean Sampling Day (Kopf *et al.*, 2015), were exploited to uncover the eukaryotic plankton biogeography (Richter *et al.*, 2019; Leconte *et al.*, 2020), taxonomy (Obiol *et al.*, 2020), and functional diversity (Delmont *et al.*, 2020). Metagenomic data often do not contain a sufficient amount of data to reconstruct nuclear genomes, but mitochondrial and plastid genomes, owing to their smaller size and a higher number of copies, may be potentially reconstructed from those data. Most often the mitochondrial genomes (Andújar *et al.*, 2015; Crampton-Platt *et al.*, 2016) or only single genes, such as 16S rDNA (Piganeau and Moreau, 2007; Piganeau *et al.*, 2008), are reconstructed from the metagenomic data. Organellar genomes have been shown to provide suitable data to address questions about microbial eukaryotes’ evolution and ecology (Cuvelier *et al.*, 2010; Kim *et al.*, 2011; Wideman *et al.*, 2020). However, organellar data are mostly unexplored in the metagenomes, since they are often classified as bacterial sequences, and are thus removed from the eukaryotic genome assemblies (Delmont *et al.*, 2020; Duncan *et al.*, 2020).

Only a few approaches dedicated to the processing of the eukaryotic fraction from the metagenomic data exist. They might be split into those developed to analyse raw reads (Wood *et al.*, 2019) or single genes (Schön *et al.*, 2020), both of which strongly depends on the reference databases. Alternatively, the eukaryotic nuclear genomes might be reconstructed from the data using one of the two main existing pipelines. In the first approach, the assembled contigs are binned, visualized, and manually refined using Anvio’O (Eren *et al.*, 2015; Delmont and Eren, 2016), whereas the second approach, used in EukRep, assumes initial separation of contigs into two domains (Prokarya and Eukarya), and then binning within those two groups independently (West *et al.*, 2018). Both approaches have been successfully used for obtaining partial nuclear eukaryotic genomes but failed to correctly classify the organellar fraction in the metagenomic data (Duncan *et al.*, 2020; Delmont *et al.*, 2020). Only one tool, MitoZ, was designed explicitly for the organellar data, but it is only applicable for the assembly, identification and analysis of the animals’ mitochondrial genomes (Meng *et al.*, 2019).

The most widely used tools for biological sequence comparison are alignment-based methods such as Smith-Waterman algorithm (Smith and Waterman, 1981) and its further developments such as BLAST (Altschul *et al.*, 1990) or BLAT (Kent, 2002). Several binning algorithms relying on the alignment-based approach, such as *taxator-tk* (Dröge *et al.*, 2015), have been proposed for the taxonomic assignment of DNA sequences in metagenomes. Although the alignment-based methods are the most accurate for sequence comparisons, they fail if sequences are highly divergent or the reference database is limited. These methods are also computationally intensive for large datasets (Ren *et al.*, 2018). For those reasons, the use of alignment-based methods for metagenomic data is relatively confined. Alignment-free methods, based on *k*-mers or DNA substrings, provide promising alternatives to overcome the weaknesses of alignment-based methods (Ren *et al.*, 2018). The usage of alignment-free methods is currently rapidly growing, and especially machine learning approaches have been used extensively for classification of various types of sequences from metagenomes (Liang *et al.*, 2020; Krawczyk *et al.*, 2018). The most potent approaches are based on deep learning, a family of machine learning methods based on artificial neural networks. Those methods best exploit large and multidimensional data sets and tackle intricate patterns in the data (Angermueller *et al.*, 2016).

The most broadly used tool for the eukaryotic metagenomics is EukRep, which uses *k*-mer frequencies and linear SVMs for DNA sequences classification (West *et al.*, 2018). It was shown to be useful for obtaining high quality nuclear eukaryotic genomes from complex environmental samples (West *et al.*, 2018), but lacks features which would enable proper organellar genomes classification. Here, we introduce Tiara, a deep-learning-based approach for identification of eukaryotic sequences in the metagenomic data sets. Its two-step classification process enables to classify nuclear and organellar eukaryotic fractions and subsequently divide organellar data into plastid and mitochondrial. Tiara outperforms EukRep in terms of prediction accuracy and calculation time.

## 2. Methods

In the first step, Tiara classifies assembled DNA sequences (contigs) into six classes: Archaea, Bacteria, Prokarya (either Bacteria or Archaea but with no specific classification), Eukarya, organelles, and unknowns (Fig. 1, the first step of classification). In the subsequent step, the organellar sequences are further classified into plastids, mitochondria, and unknowns (Fig. 1, the second step of classification). For the classification, the method employs two feed-forward neural networks and *k*-mer compositions.

**Figure. 1.**
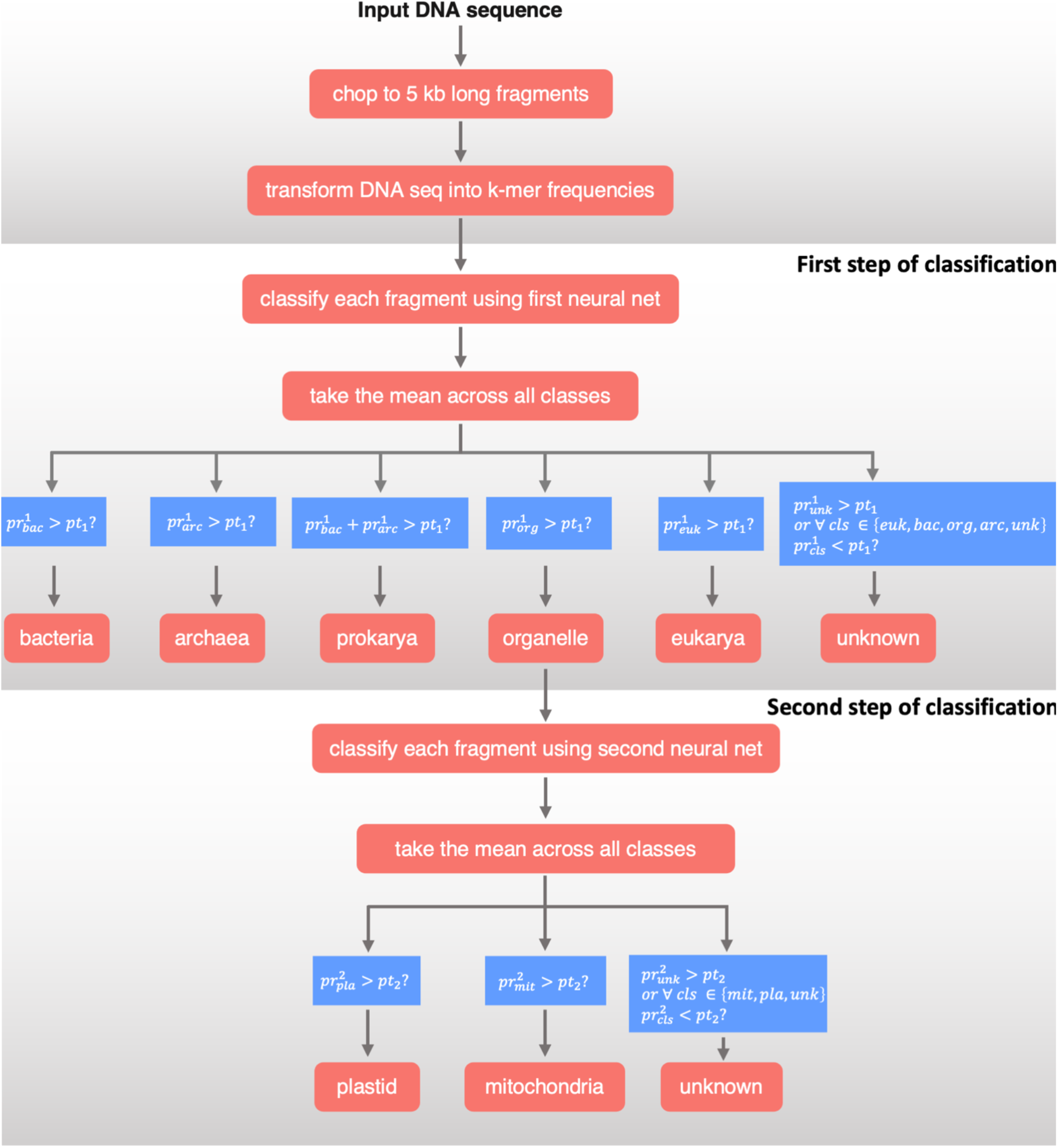
Scheme of the main steps of dataflow implemented in Tiara.

### 2.1 Training and test datasets

We decided to take advantage of taxonomic information to emerge well balanced and not overlapping train and test datasets. We independently picked genomes for each category to obtain a diverse final dataset despite differences in genomes’ lengths and their representation in reference databases.

Training datasets were prepared based on 8,220 genomic sequences (Supplementary Table S1) from three domains of life: Eukarya (4381) [nuclear (73), plastid (2260), and mitochondrial genomes (2048)], Bacteria (1860) and Archaea (1979). Data were downloaded from the NCBI Genome database (Agarwala *et al.*, 2018) and the Joint Genome Institute (JGI) (Grigoriev *et al.*, 2012). Eukaryotic nuclear genomes were chosen manually and included genomes from large groups of eukaryotes representing all supergroups, i.e., Stramenopiles, Alveolates, Chlorophyts, Rhodophytes, Amoebozoa, Opisthokonta and Discoba). Mitochondrial genomes from the NCBI genome database marked as: “Fungi”, “Plants”, “Protist” and “Other” were selected. In the case of animal mitochondrial genomes, we selected only those marked as “Other Animals” and “Insects” as representatives of this large group of small and homogeneous genomes. For plastid representation, we downloaded all available plastid genomes annotated as “Green algae”, “Protist” and “Other” in NCBI, and one representative per each genus of Land Plants to avoid overrepresentation of plants’ plastid genomes. In the case of bacterial genomes, we selected one representative from each genus present in the NCBI Genome with the best assembly quality. Due to the insufficient number of complete archaeal genomes and a dominant number of low-quality genomes derived from metagenomic initiatives, we used the best quality genomes for each archaeal species present in the NCBI genome database. Subsequently, all genome sequences were split into 5 kb fragments, and for bacterial genomes, 10% of fragments of each genome were randomly picked to achieve the better class balance between eukaryotes and prokaryotes. Fragments containing other letters than {A, T, G, C} were filtered out. Subsequently, we selected a subset of the training set as a test set, and those genomes were removed from the training set.

To test Tiara and compare it with EukRep, we prepared a test set based on 400 genomic sequences. Genomes selected as test set are not present in EukRep training dataset nor the Tiara training set and were selected with maximum overlap with the training genomes set at the genus level. The test set contains 165 eukaryotic genomes (105 nuclear, 28 plastidial and 32 mitochondrial) and 385 prokaryotic genomes (306 Bacteria and 79 Archaea). We supplemented the test dataset with data from large eukaryotic and prokaryotic groups (such as Cryptophyta, Haptophyta and Glaucocystophyta in case of eukaryotes and Candidate Phyla Radiation group, in case of prokaryotes) not represented in training set to test the Tiara ability of proper classification of divergent genomic sequences. Genomes were downloaded from NCBI and JGI databases (Supplementary Table S2). The genomes with less than 20 contigs were chopped into 100 kb long chunks, and less contiguous assemblies remained unchanged to reflect the condition of metagenomic assemblies. Mitochondrial and plastidial genomes were fragmented into pieces in a range of 1 kb – 75 kb (Supplementary Methods).

### 2.2 Sequence representation

We used tf-idf (tf – *term frequency,* idf – *inverse document frequency*) – a technique used in information retrieval – to represent DNA sequences. Each sequence was represented as a real-valued vector of length 4^*k*^, where *k* was the *k*-mer length. Given a set of sequences *S* and *s* ∈ *S*, we defined *tf_s_* as the oligonucleotide frequency vector for a sequence s and *idf_s_* as a vector describing the inverse document frequency of each *k*-mer:

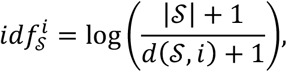

where *d*(*S, i*) is equal to the number of sequences *s*∈*S* that contain *i*-th *k*-mer (in the lexicographic order). Then the representation of the sequence s was calculated as: *v_s_* = *tf_s_⊗idf_s_*, where ⊗ is the piecewise multiplication of vectors. The vectors are then normalized to sum to one. The effect of this representation is that k-mers that occur in many DNA fragments weigh less to the prediction compared to k-mers present in only a few DNA fragments.

We have written our version of oligonucleotide frequency (*tf_s_*) calculation and an *idf_s_* vector calculation method that works online (processing one sequence at a time).

### 2.3 Classification system

We employed a two-stage classification method. In the first stage, the input sequences are classified into six classes: bacteria, archaea, prokaryote, eukaryote, organelle or unknown. The second stage differentiates between organelle subclasses: mitochondria, plastids, and unknown (Figure 1). This two-stage process relies on the two distinct two-layer feed-forward neural network architectures. Hyperparameter selection and training procedure are described in subsection 2.4.

During the classification process, we split the sequences into smaller fragments (5 kb). We then classify each fragment separately and take the mean probability for each class, resulting in five (in the first step of classification) or three (in the second step) values. We use 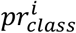 notation to describe the mean output of the neural network at stage *i* for a specific class. The classification is performed based on the probability thresholds *pt_i_*, *i* ∈ {1,2}, one for each classification stage. The sequence at stage ¿ is classified to a class if 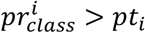. The exception from this rule is the situation where 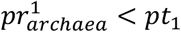 and 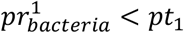, but 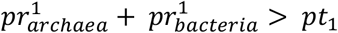. The sequence is then classified to a more general prokaryota class. If none of the above conditions are met, the sequence is classified to an unknown class.

### 2.4 Neural network architectures

#### Training

We implemented and trained our models using PyTorch (Paszke *et al.*, 2019) and skorch (Tietz *et al.*, 2017) packages using negative log-likelihood loss and an Adam optimizer (Kingma and Ba, 2015). We split the data into training (90%) and validation (10%) sets using a stratified splitting strategy: in each set, the proportion of sequences from each class was the same. The batch size was set to 128. Validation dataset was used to determine the best model.

#### Choosing the best models

To choose the best architectures we performed a search over several hyperparameters: lengths of k-mers, number of nodes in the first and second neural network layer, dropout probability, learning rate and the number of epochs. We used several metrics to compare the models: accuracy, mean precision, mean recall and mean F1 score (Supplementary Methods). The means were taken across all classes. We evaluated the models on a validation set with a probability threshold of 0.5. For consistency, we picked the architecture with the highest average mean F1 score across all learning epochs. The results of the search are in Supplementary Tables S3 and S4. To choose the architectures for the first stage, 15,900 hyperparameter combinations were tested, whereas for the second stage 17,600 combinations were tested.

### 2.5 Implementation and availability

We developed our tool in Python 3.8, with use of the libraries skorch (Tietz *et al.*, 2017), PyTorch (Paszke *et al.*, 2019), biopython (Cock *et al.*, 2009), numba (Lam *et al.*, 2015), joblib (Varoquaux and Grisel, 2009) and tqdm. Tiara code is freely available under MIT license, and the code is stored on GitHub (https://github.com/ibe-uw/tiara). The models used in the program by default are the best models for each class (marked in bold in Supplementary Table S6), but the user can choose other optimal models for each *k*-mer length. By default, Tiara returns tabular output with a class assigned to each contig name but optionally allowing the user to output classified sequences to separate files in fasta format.

## 3. Results

We implemented in Tiara a two-stage approach for classification of eukaryotic sequences from assembled metagenomic data. In the first stage, Tiara classifies sequences into six categories (Archaea, Bacteria, Prokarya, Eukaryota, organelles or unknown). In the second stage, putative organellar sequences are classified into plastids and mitochondria or unknown. Each stage of classification encapsulates trained neural network model. Next, we evaluated its performance and compared with EukRep using independent test dataset and showed usability using real metagenomic data.

### 3.1 Performance comparison of different *k*-mer sizes

We searched hyperparameter space to obtain the best neural network models (nearly 35000 models) using a validation dataset. The best hyperparameters for each *k*-mer length and classification stage for an optimal number of epochs are shown in Supplementary Table S5. The best first stage neural network was trained with *k*-mer length 6, and had two layers with 2048 and 1024 nodes, respectively. The best neural network in the second stage of classification used a *k*-mer length 7 and had two layers with 128 and 64 nodes. For both stages, we used the dropout probability of 0.2. The first best model was trained for 41 epochs using a learning rate equal to 0.001, and the second for 47 epochs with a learning rate of 0.01. The comparison of the best models for each *k*-mer length (in bold) with a sub-optimal architecture (both layer sizes equal to 32, learning rate of 0.01 and 0.5 dropout probability – in italics) shows that larger neural networks are necessary to identify the biological signal present in the DNA sequences (Supplementary Table S5).

### 3.2. Performance of trained models

#### Classification of eukaryotic and prokaryotic sequences

Depending on the *k*-mer length, Tiara achieved on the test dataset mean prediction accuracy between 98.65% and 98.93% for prokaryotic genomes and between 95.94% and 98.83% for the nuclear genomes (Table 1). The best ratio between prediction accuracies for each class was noted for *k*-mer 6. Using this model, 96% of nuclear eukaryotic and 98% of prokaryotic genomes were classified with higher or equal prediction accuracy to 90%. Only three prokaryotic genomes have been classified with accuracy lower than 50%. However, most of the contigs derived from these genomes were assigned as “unknown”, and only two were classified as a eukaryote. All of these genomes were small and highly reduced, and they belonged to symbionts or parasites. This bias was also previously observed for EukRep (West *et al.*, 2018). We checked probability outputs for them and observed strong organellar signal for two endosymbionts, which might reflect the reductive evolution of their genomes.

**Table 1.** Comparison of accuracy for Tiara and Eukrep tools. All Tiara tests have been done by default with 0.65 probability cutoffs. EukRep was tested with default settings. The best model for a given class is shown in bold.

| Software | $k$-mer | Avg. accuracy Eukarya<br>(n=105) | Avg. accuracy<br>Prokarya (n=385) | Avg. accuracy<br>organelles(n=60)* |
| --- | --- | --- | --- | --- |
| Tiara | 4 | 0.9593 | 0.9871 | 0.9411<sup>mt</sup>/0.9708<sup>pt</sup> |
| Tiara | 5 | 0.9641 | 0.9865 | 0.9841<sup>mt</sup>/0.9981<sup>pt</sup> |
| Tiara | 6 | **0.9883** | **0.9893** | **0.9886<sup>mt</sup>/0.996<sup>pt</sup>** |
| EukRep | 5 | 0.963 | 0.9849 | 0.4738<sup>mt</sup>/0.2979<sup>pt</sup> |

\* In the case of EukRep we calculated the ratio of organellar fragments classified as Eukaryote.

Importantly, Tiara achieved high accuracies (above 90%) for genomes from groups of taxa that were absent in the training dataset like eukaryotic haptophytes and cryptophytes or prokaryotic CPR, which indicates that our models are not overfitting despite employing complex neural networks. Hence, Tiara will be able to classify contigs of novel evolutionary lineages correctly.

#### Classification of organellar sequences

Tiara achieved high prediction accuracy for a testing set of organellar genomes (28 plastidial and 32 mitochondrial) with an average accuracy above 95% (Table 1). The best average accuracies were observed for *k*-mer 6 (pt: 99.60%; mt: 98.86%). Similar to the nuclear genomes’ classification, the accuracy of organellar genomes classification increased with *k*-mer length. (Supplementary Table S6). Analysis of fragmented organellar genomes (mitochondria: (1-5 kb) and plastids (1 kb-75 kb) for *k*-mers *k* = {4-6}) showed that prediction accuracy increased with the fragment length (Supplementary Figure S1 and S2;). For organellar sequences longer than 3 kb, accuracy was higher than 90%, and for sequences longer or equal to 5 kb – close to 100%. In the second stage, most of the sequences were assigned correctly to a given class (higher than 90%) if the sequence was longer than 3 kb.

### 3.3 Performance comparison between Tiara and EukRep

We compared Tiara with EukRep – a tool designed for the classification of eukaryotic and prokaryotic sequences from metagenomic data (West *et al.*, 2018). EukRep was previously shown to outperform alignment-based methods. Thus, we compared Tiara only with EukRep, which is currently the state-of-the-art method. Finally, it enabled the fast identification of eukaryotic contigs and further forming eukaryotic MAGs (Metagenome Assembled Genomes). EukRep, similarly to Tiara, transforms DNA sequences into *k*-mer frequencies, but then uses linear-SVM (implemented in scikit-learn) for predictions, whereas Tiara uses sequential feed-forward neural networks. EukRep, as a binary classifier, separates data only into two classes (domains): eukaryotes and prokaryotes. Therefore, EukRep hasn’t been trained on organellar DNA, so it was unclear how it classifies those sequences.

Tiara scored slightly better than EukRep, with the prediction accuracy of eukaryotic genomes 2.53% higher and prokaryotic genomes – 0.44% higher (Supplementary Table S2). We also calculated the difference in prediction accuracy between Tiara and EukRep for each genomes pair (Fig. 2) to examine prediction accuracy in details. For eukaryotic genomes (105), Tiara got better accuracies in 63 cases, and only for 25 genomes, the results were worse. Whereas prokaryotes were classified more evenly and for 275 genomes, both tools have the same results; in 89 cases, Tiara was better than EukRep, and in 21 cases it was worse.

**Figure. 2.**
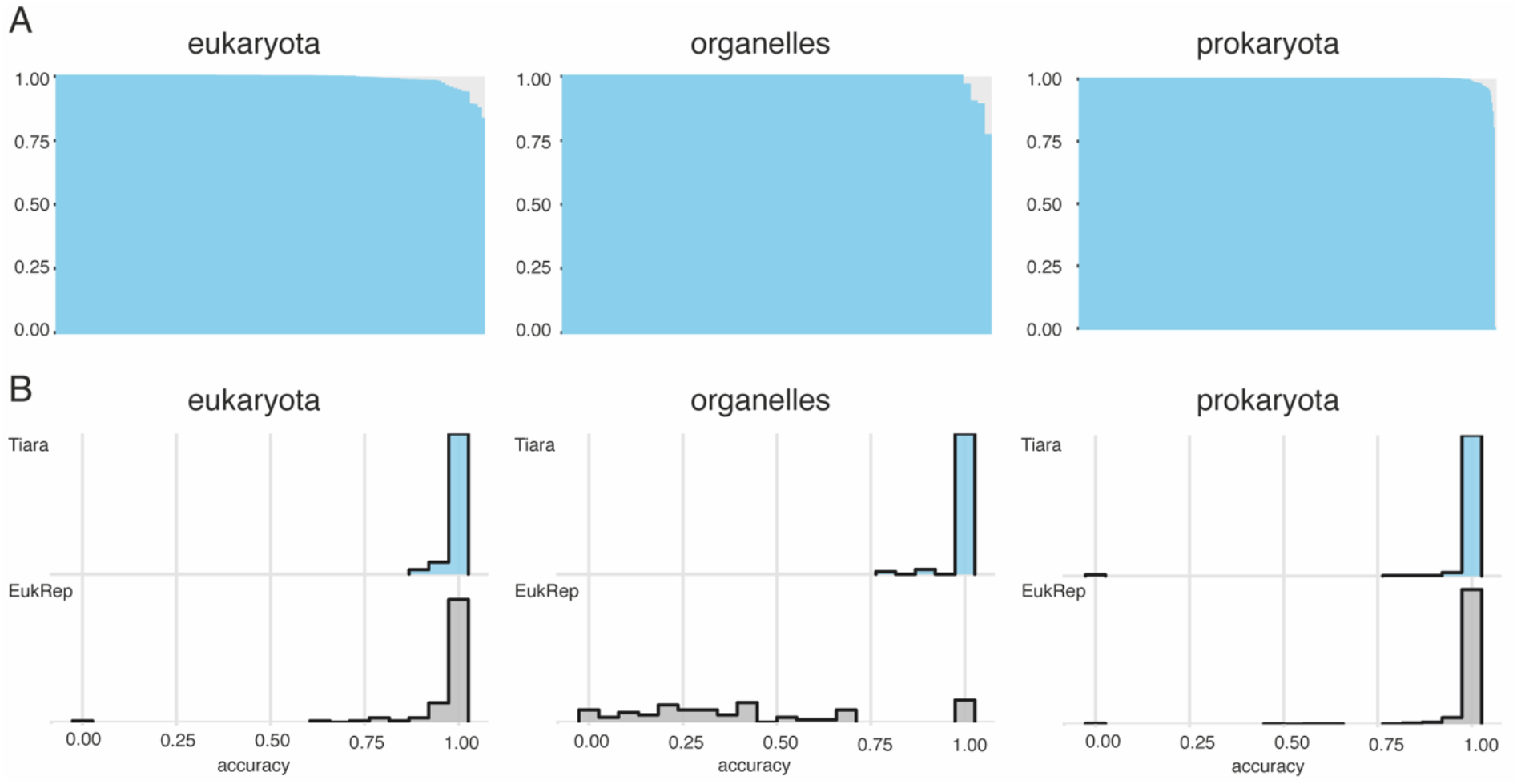
Efficiency of Tiara classification and comparison to EukRep using a set of test genomes. Test genomes were divided into three groups: eukaryotic nuclear genomes (eukaryota), plastid and mitochondrial genomes (organelles), and archaeal and bacterial genomes (prokaryota). (A) Accuracy of Tiara for each genome in three groups (B) Histogram of the density of accuracy for EukRep and Tiara for three groups of genomes. All tests have been performed using the model with *k*-mer 6 and 0.65 probability cutoff.

To test the organellar genomes classification by EukRep, we used a set of fragmented plastid (10 kb) and mitochondrial (5 kb) genomes. EukRep assigned only 47% of mitochondrial and 30% of plastid contigs as eukaryotic sequences (Supplementary Table S2).

Additionally, we checked the speed performance of both approaches (Supplementary Methods). Tiara supports parallel execution, whereas EukRep uses all cores available. However, for one core, Tiara was two times faster than EukRep. Finally, Tiara using 12 cores classified the test genome roughly five times faster than EukRep and reached a speed of 6.8 Mbp per second. (Supplementary Table S7).

### 3.4 Classification of sequences from the real data

To examine our approach on real metagenomic data, we used three datasets from the Tara Oceans project (Pesant *et al.*, 2015). We selected three samples from the Mediterranean Sea (ERR1726574, ERR1726673, ERR868402) from the same station, representing three different size fractions associated with protists (Supplementary Methods, Supplementary Table S8). Data were assembled (Supplementary Methods; Table S8) and used for further analyses. We tested Tiara with three *k*-mer lengths (4, 5, and 6) and three minimum sequence lengths (1000, 3000, 5000 bp) for the first stage of classification (Supplementary Methods).

#### Composition of metagenomic samples

In the metagenome of the smallest size fraction (0.8-20 μm), prokaryotes seemed to prevail, as the majority of contigs were classified as Bacteria, Archaea or Prokarya (up to 97% for k=5); however, datasets from larger size fractions (20-180 and 180-2000 μm) were dominated by eukaryotes (up to 96% for k=6) (Supplementary Figure S3 and Table S9). Contribution of contigs assigned to eukaryotes was the highest using model with the *k*-mer length k=6, which is in line with the results obtained from the test datasets. The organellar fraction’s overall contribution was low in assembled data and ranged between 0.26% and 4.2% across datasets and analysis variants. Nevertheless, organellar contigs were among the longest ones and exceeded 50 kb for sample ERR1726673 (Supplementary Table S10).

#### Analysis of large fragments classified as organelles

We annotated 21 contigs assigned as organellar and longer than 10 kb (Supplementary Methods). Among those 21 contigs, 13 were annotated as mitochondrial and seven as plastidial, and for one, blastN reported no significant hits. (Supplementary Table S10). For the smallest size fraction (0.5-20 μm), all five analyzed contigs were derived from two plastid genomes, belonging to the dictyochophycean *Florenciella parvula* and the green alga *Pycnococcus provasoli.* Three fragments of the *Pycnococcus* plastid genome together accounted for the 58.2% of its estimated size and carried 38 genes. The largest taxonomic diversity of contigs was detected in the size fraction 20-180 μm; organellar genomes of nine protists (diatoms, ciliates) and animals (crustaceans, insects and molluscs) were identified. For the largest size fraction (180-2000 μm), we identified three partial mitochondrial genomes that belonged to animals (crustaceans and hydrozoans).

## 5. Discussion

We developed Tiara, a machine learning-based tool, which can efficiently and accurately separate eukaryotic sequences from the prokaryotic ones to overcome difficulties with eukaryotic data classification in the metagenomic data. Tiara doesn’t rely on large reference databases and can be efficiently utilized in pipelines for identifying eukaryotic scaffolds and binning into MAGs. Tiara is also the first tool designed to consider organellar sequences as a separate class, allowing further analyses.

Trained models encapsulated within Tiara scored high accuracy for validation and test dataset, suggesting that models are not overfitting. Moreover, longer *k*-mers coupled with large networks resulted in the best performance in both classification stages, confirming that complex neural networks can better identify informative signal within DNA sequences.

Using a test dataset, we have shown that Tiara performs similarly to EukRep (representing the current state-of-the-art) in terms of prokaryotes classification, and outperformed it in terms of classification of eukaryotes with considerably lower calculation time. Tiara was trained on a much larger data set than EukRep and employed neural networks, which allowed longer *k*-mer (6) usage and performed better with more complex data. EukRep uses linear-SVMs, which are less effective when dealing with multidimensional data. Crucially, Tiara correctly classifies sequences from organellar genomes (up to nearly 100% of plastid sequences and 99% of mitochondrial sequences). In contrast, EukRep recovered an only small portion of organellar fragments (approximately 33%) classifying them as eukaryotic ones (Table 1).

Analysis of the metagenomic data from the Mediterranean Sea allowed the classification and reconstruction of the organellar genomes. Tiara classified most of the contigs as nuclear genomes for the fraction larger than 20 μm, and the smallest fraction was dominated by prokaryotes, as already reported in previous studies (Tully et al. 2018). Analysis of a range of *k*-mers and a sequence length cut-off confirmed that the lowest false positive rate is achieved for *k*-mer 6 with a minimum sequence length of 3 kb. Even though organellar fragments constituted less than 10% of contigs, they were among the longest ones. Thanks to Tiara, we reconstructed three partially complete plastid genomes and twelve almost complete mitochondrial genomes from the Mediterranean Sea data set. Only one plastid genome was classified to the species level (99% of identity); however, it most likely represents a different strain than those deposited as reference data in databases. For all mitochondrial genomes, the classification was restricted by the NCBI database’s lack of close reference. Six of the identified genomes had only a moderate similarity to crustacean genomes of *Undinula vulgaris* (~72%) and *Paracyclopina nana* (~76%). This result suggests that we recovered mitochondrial genomes of crustacean species currently not represented in the NCBI database.

Still, some challenges remain. The classification of shorter sequences might be wrong because those sequences are less informative. Thus, we recommend analyzing sequences that are longer than 3 kb to reduce the false-positive rate. The classification of eukaryotic sequences might also be disturbed by the existence of NUMTs and NUPTs – fragments of mitochondrial or plastid genomes localized in nuclear genomes (Kim and Lee, 2018). Those fragments might be misassigned as organellar sequences. Another problem constitutes introns and other extremely divergent non-coding regions, which might significantly disturb *k*-mer frequencies locally, resulting in the wrong prediction if a given sequence is too short to retain distinctive signal. Finally, the relatively high similarity of rDNA operons between groups can result in misclassification of those regions. Thus, we suggest using additional tools like Phyloflash to analyze those regions (Gruber-Vodicka *et al.*, 2020).

In the current analysis, the class “unknown” from the first stage of classification has been added to the class “eukaryotic” to maximize eukaryotic sequences’ recovery. This decision was based on the assumption that ambiguous predictions will less likely fall into the class prokaryote since the set of prokaryotic genomes used for training was extensive, diverse and evenly-sampled. Consequently, there is a high chance that those fragments belong to eukaryotic genomes. The prokaryotic and viral sequences which might end up in the class “unknown” can also be easily removed during preprocessing step like binning and bin refinement. However, our assumption might slightly increase the number of false positives in the first stage of classification.

Currently, sequences shorter than the given threshold remain unclassified and should be treated separately using gene-centric approaches. Those sequences are less informative and might significantly increase the number of false positives. However, it is still worth analyzing them to detect mitochondrial and plastid genomes of rare protists.

Despite its advantages, organellar genomes so far have not been widely used, compared to metabarcoding or single-cell approaches. We hope that Tiara will enable researchers to make more use of metagenomic data. Organellar data can be employed for phylogenomic reconstruction and uncover new eukaryotic lineages, as already have been shown for mitochondrial genomes of marine heterotrophic protists (Wideman *et al.*, 2020). Organellar sequences, similarly to barcodes, are also applicable for diversity assessment and biogeographic studies. Even partial organellar genomes might be successfully used to study organellar genomes’ structure and content (Cuvelier *et al.*, 2010; Hovde *et al.*, 2014). Ultimately, all these approaches enable a deeper understanding of diversity and evolution of eukaryotes.

The reconstructed underrepresented genomes can be used to supplement existing databases. That would further reduce the false positives and allow for more precise classification. Tiara could also be broadly used for metagenomic data preprocessing to remove eukaryotic contamination, including more difficult to distinguish from prokaryotic data organellar sequences.

Our analyses have shown that publicly available metagenomic data contain a considerable number of organellar DNA fragments. Among analyzed datasets, we were able to identify organellar sequences of previously unreported plastid and mitochondrial genomes. Our results also suggest that even from low-quality data, which are insufficient for nuclear genomes assembly, almost complete organellar genomes can be reconstructed.

## Acknowledgements

We thank Stanisław Dunin-Horkawicz and Kacper Maciszewski for critical reading of the manuscript and many colleagues for carrying out beta tests of the software.

