## Supplementary materials for "Tiara: Deep learning-based classification system for eukaryotic sequences"

### Supplementary methods

#### 1. Preparation of organellar test genomes

Organellar genomes were chopped into chunks from 1000 bp up to the length of the shortest genome from a given set. Plastid genomes were fragmented into pieces of lengths = {1000, 2000, 3000, 5000, 10000, 50000, 75000}. While smaller mitochondrial genomes were fragmented into chunks of lengths = {1000, 2000, 3000, 5000}.

#### 2. Metrics definitions

Below we present definitions of metrics used to estimate the effectiveness of our models.

$$Accuracy = \frac{(TP + TN)}{(TP\ +\ TN\ +\ FP\ +\ FN)}$$

$$Precision = \frac{TP}{(TP + FN)}$$

$$Recall = \frac{TP}{(TP + FN)}$$

Where: TP = True positive; FP = False positive; TN = True negative; FN = False negative

*F1 score*

$$F1\ score = 2 * \frac{precision * recall}{(precision + recall)}$$

For tests on independent datasets (section 2.1; Supplementary Table S2) the accuracy was defined as s a sum of sequences length in a positive class divided by the sum of lengths of all classified sequences. The class eukarya is defined as a sum of sequences assigned as "organelles", eukaryotic nuclear ("eukarya") and "unknown", while prokarya is defined as the sum of sequences assigned as "prokarya", "archaea" and "bacteria”. The positive class for organellar genomes was assigned as “organelles”.

#### 3. Speed comparison of Tiara and EukRep

Tiara and EukRep were tested on the nuclear genome of a diatom *Minidiscus trioculatus* (https://mycocosm.jgi.doe.gov) in range of cores (from 1 to 12) with a minimum sequence length of 3000 bp. Since EukRep uses all available cores by the default, we measured calculation time for 12 cores available on the tested machine (Supplementary Table S7). All tests have been performed on the machine with Intel(R) Xeon(R) CPU E5-2620 v4 @ 2.10GH and, 128 GB of RAM. The workstation was running under Ubuntu Linux 16.04.

#### 4. Analysis of metagenomes from the Mediterranean Sea (Tara Oceans)

Three metagenomic datasets from one spot on the Mediterranean Sea acquired from the Tara Oceans Initiative (Pesant *et al.*, 2015) were processed. Raw reads were downloaded from NCBI Short Read Archive (accessions: ERR 1726574, ERR 1726673, ERR 868402) using SRA toolkit. Metadata associated with these datasets were stored in the Supplementary

Table S8. Raw reads have been quality checked using fastqc (Andrews *et al.,* 2010) and assembled using MEGAHIT (Li *et al.*, 2015) with default parameters. Basic assembly statistics have been calculated using QUAST (Gurevich *et al.*, 2013). Assembled contigs (Supplementary Table S8) from each dataset were analyzed using Tiara with three kmers = {4,5,6}, and three different minimum sequence lengths = {1000, 2000, 3000} (Supplementary Table S9). For taxonomic annotation of contigs longer than 10 kb assigned as organelles by Tiara using model with k=6, we used blastN against the NCBI nt database (Johnson *et al.*, 2008). Then we estimated completeness of the putative genome as a length of fragment divided by the closest blast hit representing a complete or partially complete genome. After that open reading frames (ORFs) were predicted using Prodigal (Hyatt *et al.*, 2010) in "meta" mode (Supplementary Table S10).

### Supplementary Figures

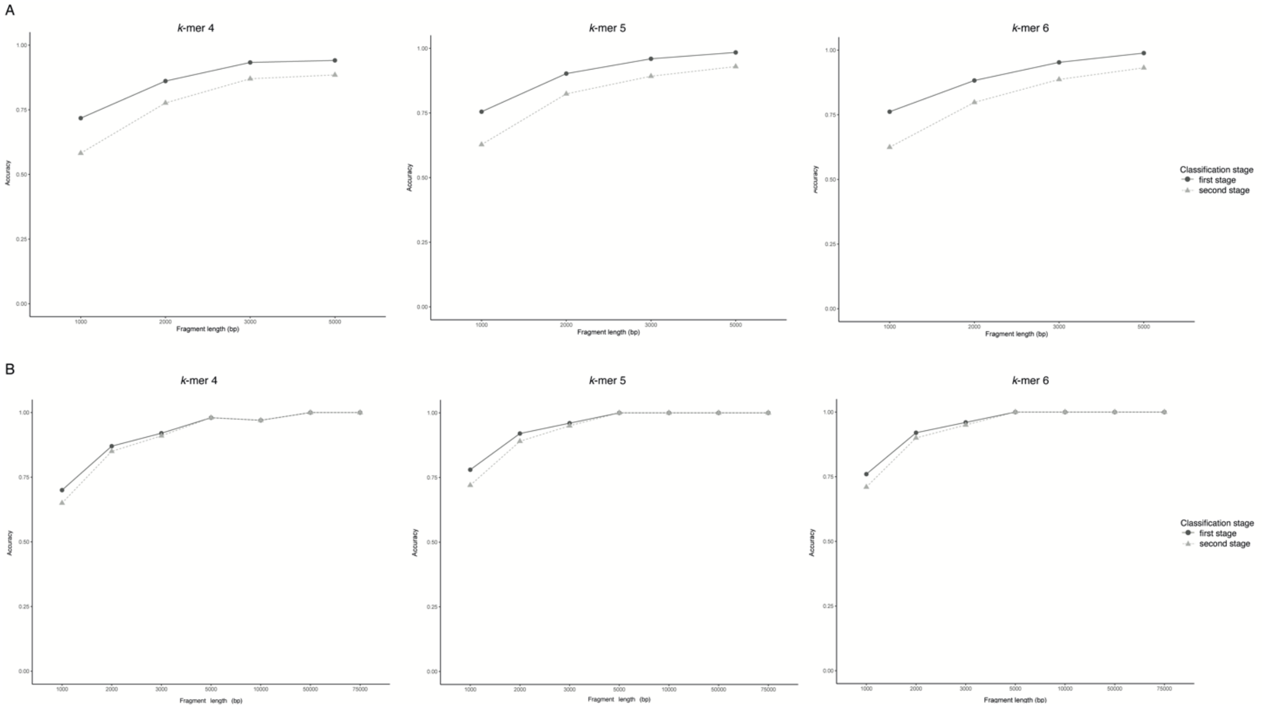


**Supplementary Figure S1.** The efficiency of the Tiara classification on test organellar genomes (mitochondrial and plastid) depending on the fragments' lengths. Tests were performed for three *k*-mers (4, 5, 6) for the first stage of classification.

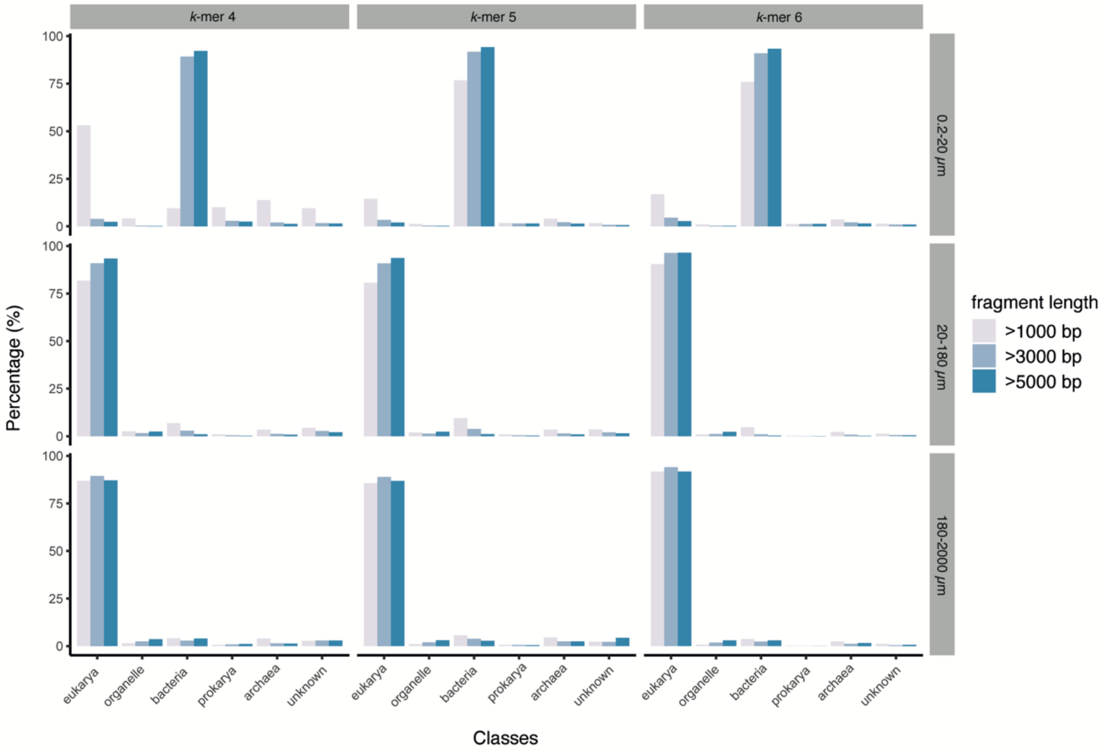


**Supplementary Figure S2.** Results of the first stage of classification of metagenomic data by Tiara. Three models with *k*-mer lengths = {4,5,6} were used, three minimum sequence lengths and three fraction sizes.

**Supplementary Tables**

**Supplementary Table S1.** List of genomes used for training models.

**Supplementary Table S2.** List of genomes used for testing Tiara and EukRep. Both software classified only sequences longer than 3000 bp. Tiara was tested using k-mers of k = {4,5,6} with probability cut-offs 0.65. EukRep was tested in recommended balanced mode.

**Supplementary Table S3.** List of hyperparameters and evaluation of 15900 searched models for the first stage of classification. Models were evaluated using validation dataset.

**Supplementary Table S4.** List of hyperparameters and evaluation of 17600 searched models for the second stage of classification. Models were evaluated using validation dataset.

**Supplementary Table S5.** Evaluation of the best models and its hyperparameters for each stage of classification with different k-mer lengths for an optimal number of epochs. The best models for each stage were marked in bold. The row presented in italics shows the sub-optimal model for the first stage.

**Supplementary Table S6.** The efficiency of the classification of test dataset of organelles depending on fragment lengths and used models. Plastid fragments ranged from 1000 bp to 75000 bp, whereas mitochondrial from 1000 bp to 5000 bp.

**Supplementary Table S7.** The calculation time of Tiara and EukRep depending on the number of cores. For test nuclear genome of *Minidiscus trioculatus* (76.188 MB) was used.

**Supplementary Table S8.** Metadata and assembly statistics for three metagenomic datasets from the Mediterranean Sea (TARA Oceans) used as a case study.

**Supplementary Table S9.** Results of the first stage of classification of metagenomic datasets by Tiara. Three *k*-mer lengths and three minimum lengths of contigs were used.

**Supplementary Table S10.** Annotation of contigs longer than 10 kb classified by Tiara as an organelle.
